# Structure and Function of the Retina of Low-density Lipoprotein Receptor-related Protein 5 (Lrp5)-deficient Rats

**DOI:** 10.1101/2021.11.05.467506

**Authors:** John L. Ubels, Cheng-Mao Lin, David A. Antonetti, Monica Diaz-Coranguez, Cassandra R. Diegel, Bart O. Williams

## Abstract

Loss-of-function mutations in the Wnt co-receptor, low-density lipoprotein receptor-related protein 5 (LRP5), result in familial exudative vitreoretinopathy (FEVR), osteoporosis-pseudoglioma syndrome (OPPG), and Norrie disease. CRISPR/Cas9 gene editing was used to produce rat strains deficient in Lrp5. The purpose of this study was to validate this rat model for studies of hypovascular, exudative retinopathies. The retinal vasculature of wildtype and Lrp5 knockout rats was stained with *Giffonia simplifolia* isolectin B_4_ and imaged by fluorescence microscopy. Effects on retinal structure were investigated by histology. The integrity of the blood-retina barrier was analyzed by staining for claudin-5 and measurement of permeability to Evans blue dye. Retinas were imaged by fundus photography and SD-OCT, and electroretinograms were recorded. Lrp5 gene deletion led to sparse superficial retinal capillaries and loss of the deep and intermediate plexuses. Autofluorescent exudates were observed, correlated with absence of claudin-5 expression in superficial vessels and increased Evans blue permeability. OCT images show pathology similar to OCT of humans with FEVR, and retinal thickness is reduced by 50% compared to wild-type rats. Histology and OCT reveal that photoreceptor and outer plexiform layers are absent. The retina failed to demonstrate an ERG response. CRISPR/Cas9 gene-editing produced a predictable rat Lrp5 knockout model with extensive defects in the retinal vascular and neural structure and function. This rat model should be useful for studies of exudative retinal vascular diseases involving the Wnt and norrin pathways.

## 1. Introduction

Canonical Wnt signaling is required for development of the retinal vasculature and the establishment and maintenance of the blood-retinal barrier (BRB) (Langen et al., 2019; Wang et al., 2012; Zhou et al., 2014). In the brain and retina, Wnt ligands, secreted by glia, bind to a frizzled (Fzd) family member receptor complex in endothelial cells. Canonical signaling through inhibition of GSK-3α/β kinase stops β-catenin degradation by the adenomatosis polyposis coli (APC) degradation complex. β-catenin then translocates to the nucleus and contributes to the activation of genes that regulate vascular angiogenesis and the development (barriergenesis) and maintenance of the vascular endothelial barrier (Ye et al., 2009). In the retina, norrin, a Müller cell-derived ligand that binds specifically to the Fzd4 receptor complex, also activates this pathway (Ke et al., 2013).

Both Wnt- and norrin-induced signaling require the co-receptor, low-density lipoprotein receptor-related protein-5 or 6 (LRP5 or 6). LRP5 complexes with the Fzd4 receptor at the cell membrane and interacts with the ligand through its extracellular β-propeller domains, resulting in a potentiated β-catenin response (Berger et al., 1992; Diaz-Coranguez et al., 2020; Hua et al., 2018; Joiner et al., 2013; Ke et al., 2013; Ubels et al., 2020).

Mutations in members of the Fzd4 receptor complex or signaling pathway cause a spectrum of inherited exudative retinopathies that result in blindness. These diseases are characterized by hypovascularization, neovascularization due to hypoxia, fibrosis, breakdown of the BRB causing exudates, retinal traction, and detachment. Norrie disease and Coat’s disease are caused by mutations in the norrin gene (*NDP*) (Black et al., 1999; Parzefall et al., 2014), while the disorders referred to as familial exudative vitreoretinopathy (FEVR) can be caused by mutations in *LRP5*, *NDP*, *Fzd4*, *TSPAN12,* or β-catenin (Fei et al., 2014; Gilmour, 2015; Masuda et al., 2016; Nikopoulos et al., 2010; Panagiotou et al., 2017; Schatz and Khan, 2017; Warden et al., 2007; Xu et al., 2014). These genes are also mutated in 3-11% of the patients with retinopathy of prematurity (Els et al., 2010; Hiraoka et al., 2001; MacDonald et al., 2005; Shastry et al., 1997), which has pathophysiological similarities to diabetic retinopathy in adults (Kermorvant-Duchemin et al., 2010).

LRP5 is also involved in bone development, acting in the osteoblast lineage. Loss of function mutations of *LRP5* cause reduced bone mass and bone mineral density leading to osteoporosis (Cui et al., 2011; Holmen et al., 2004; Joeng et al., 2011; Maupin et al., 2013; Riddle et al., 2013). The effects of an *LRP5* mutation on retina and bone come together in the rare disease osteoporosis pseudoglioma syndrome (OPPG) (Ali et al., 2005; Gong et al., 2001; Maltese et al., 2017). Children with an *LRP5* mutation causing OPPG have severely impaired vision at birth due to retinal hypovascularization and retrolental fibrovascular tissue (pseudoglioma). They also suffer from osteoporosis, with brittle, deformed bones before the age of five.

Effects of *Lrp5* mutations on the retinal vasculature and skeletal system have been studied using mouse models developed using embryonic stem cells and standard genetic engineering techniques (Ye et al., 2009; Zhou et al., 2014; Chen et al., 2012; Huang et al., 2016; Kato et al., 2002; Wang et al., 2016; Wang et al., 2019; Xia et al., 2010). However, the small size of the bones makes the mouse model sub-optimal for orthopedic research. To take advantage of the larger skeletal size in rats, we used CRISPR/Cas9 technology to produce three rat lines with 18 bp or 22 bp deletions or an allelic inversion in the *Lrp5* gene, all resulting in loss of function mutations. As we recently reported, *Lrp5* deletion in the rat results in reduced bone mineral density and bone mass in trabecular bone and decreased cortical bone size Ubels et al., 2020).

We also investigated the effect of *Lrp*5 gene deletion on the retinal vasculature and demonstrated markedly abnormal vascularization in all three rat lines (Ubels et al., 2020). The superficial vasculature was disorganized, with small vessels nearly absent in large regions of the retina. Quantitatively, the vascularized area of the retina and the number of vessel branch points was significantly reduced. The retinas also contained extensive autofluorescent exudates, suggesting a defective BRB (Ye et al., 2009; Xia et al., 2008). Deletion of *Lrp*5 using CRISPR/Cas9 thus resulted in a model in which bone and retinal pathology can be studied in the same rats.

The initial study of this *Lrp5* knockout rat model was limited to the superficial retinal vasculature architecture. The purpose of the present study was to further investigate retinal changes in the rat model of *Lrp5* gene deletion and determine whether this model might be useful for more detailed studies of mechanisms that cause hypovascular retinopathies. Using the rat strain with the 18 bp deletion, we characterized the changes in the intermediate and deep vascular plexuses and examined the neural retina structure by histology. We evaluated the retinas of Lrp5-deficient rats in vivo using fundus photography and optical coherence tomography (OCT). The presence of exudates, as detected microscopically and by OCT, was correlated with impaired BRB function by measurement of vascular permeability to Evans blue dye (Xu et al., 2001) and by investigating the expression of claudin-5, an essential component of junctional complexes that form the BRB (Diaz-Coranguez et al., 2017). The effect of neurovascular unit pathology on visual function was examined by electroretinography.

## 2. Methods

### 2.1. Knockout of Lrp5

Maintenance of Sprague-Dawley rats (Charles River Laboratories, Wilmington, MA) and all experimental protocols were approved by the Institutional Animal Care and Use Committees of the Van Andel Institute and the University of Michigan and conformed to the ARVO Statement for the Use of Animals in Ophthalmic and Vision Research. Both male and female rats were used in the study. There was no statistical difference between the sexes in this or our previous study (Ubels et al., 2020), so all data were combined.

As previously described in detail (Ubels et al., 2020), three rat lines with deletions in exon 2 of the *Lrp5* gene were created using a modified CRISPR/Cas9 protocol. Most of the data in this report are from rats with an 18 bp in-frame deletion in exon 2 of *Lrp5*. This deletion causes loss of six amino acids in β-propeller 1 of Lrp5 which, as a result, cannot complex with Wnt.^7^ Images from rats with a 22 bp deletion in exon 2, also resulting in a defective β-propeller 1, are provided as supplementary data. The founders were backcrossed to wild-type Sprague-Dawley rats twice before intercrossing to generate animals for this study. The animals were genotyped, and the lack of *Lrp5* gene expression was assessed, as previously described (Ubels et al., 2020).

### 2.2. Staining of the Vasculature in Retinal Flat Mounts and Frozen Sections

For imaging of the superficial retinal vasculature, rats were euthanized with CO2, and the eyes were enucleated and fixed in 4% paraformaldehyde. As previously described (Chen et al., 2012; Wang et al., 2016; Ubels et al., 2020),^7, 35, 36^ retinas were incubated in 1% Triton X-100, stained with Alexa Fluor 594-conjugated *Giffonia simplifolia* isolectin B_4_ (Thermo Fisher Scientific), washed in PBS, flat-mounted on Superfrost slides, and cover-slipped in Prolong Gold antifade reagent (Thermo Fisher Scientific).

To examine the intermediate and deep vascular plexuses, eyes of 2.5 to 3 month-old rats were fixed in Davidson’s fixative (20% isopropanol, 3% glacial acetic acid, 4% formaldehyde, in DI water). The cornea and lens were removed, and the eyecups were rinsed in PBS. The eyecups were placed in 30% sucrose in PBS at 4°C overnight, embedded in OTC medium, frozen in dry ice/isopentane, and stored at −80°C. The retinas were cryosectioned at 12 μm. Sections were mounted on Superfrost slides, stained with Alexa Fluor 594-conjugated isolectin B_4_ for 5 minutes, washed with PBS, and cover-slipped in Prolong Gold antifade reagent containing DAPI (Wang et al., 2016; Ubels et al., 2020).

To detect claudin-5 in the retinal vasculature, 12 μm sections were prepared and mounted in the same manner and blocked with normal goat serum for 60 minutes. The sections were then incubated with Alexafluor 488-conjugated claudin-5 monoclonal antibody (4C3C2, Thermo Fisher Scientific) overnight at 4°C, washed with PBS, and cover-slipped in Prolong Gold antifade reagent containing DAPI.

Fluorescent images were captured using a Zeiss Axio Imager A2 microscope with ET – DAPI/FITC/Texas Red filters (Carl Zeiss AG, Oberkochen, Germany) and BIOQUANT OSTEO 2019 v19.2.60 software (BIOQUANT Image Analysis Corp., Nashville, TN).

### 2.3. Histology

Eyes of 2.5 to 3 month-old rats were enucleated and whole globes were fixed in Davidson’s fixative. Eyes in fixative were sent to Excalibur Pathology, Inc., Norman, OK, for processing, where 5 μm serial sections taken 3 mm each side of the optic nerve were mounted and stained with hematoxylin and eosin. Bright-field images were captured using a Zeiss Axio Imager A2 microscope and BIOQUANT OSTEO 2019 v19.2.60 software, which was also used to measure thickness of retinal layers.

### 2.4. Fundus Photography and Optical Coherence Tomography

Pupils of 3-month-old rats were dilated with one drop of 1% tropicamide followed by one drop of 2.5% phenylephrine, and the rats were anesthetized with Ketamine and Xylazine (90 and 10 mg/kg body weight). Bright-field fundus photographs were taken with a Phoenix Micron III Retinal Imaging System (Phoenix Technology Group, Pleasanton, CA). In vivo images of retinal anatomy were acquired using a Bioptigen Envisu Preclinical Spectral Domain Optical Coherence Imaging System (Leica Microsystems, Buffalo Grove, IL). Rectangular volumes consisted of 1000 A-scans by 100 B-scans over a 2.6 x 2.6-mm area. Images were taken centered on the optic nerve head and also temporally and nasally to the optic nerve head. Peripheral retinal thickness was quantified with the Bioptigen Diver software at 2 mm from the optic nerve head.

### 2.5. Measurement of Vascular Permeability

Because the CRISPR/Cas9 *Lrp5* knockout model was created in albino rats, it was not possible to image the retinal vasculature using fluorescein angiography. Permeability of the retinal vascular of three-month-old rats was therefore assessed using Evans blue dye (Xu et al., 2001). Briefly, rats were anesthetized with Ketamine and Xylazine, and 45 mg/kg body weight of Evans blue dye (Sigma-Aldrich, St. Louis, MO; E-2129) and saline was injected via the femoral vein. After 2 hours, 0.3 ml of blood was drawn from the vena cava to obtain a plasma sample. The rats were then euthanized and perfused with warm saline at 66 ml/minute for 2 minutes via the left ventricle. Retinas were harvested, dried overnight in a Savant SpeedVac vacuum concentrator (Thermo Fisher Scientific), and weighed. The dye was extracted from retinas with formamide (Sigma- Aldrich, St. Louis, MO) at 70°C overnight. Dye concentrations in retina extracts and plasma samples were determined by absorbance at 620 and 740 nm using a FLUOstar Omega microplate reader (BMG Labtech Inc., Cary, NC).

### 2.6. Electroretinograms

A Diagnosys Celeris System (Diagnosys, Lowell, MA) was used to record electroretinograms. Three-month-old rats were dark-adapted overnight. Topical 0.5% tropicamide was applied to dilate the pupil, and the corneas were anesthetized with 0.5% proparacaine. Rats were anesthetized with an intraperitoneal injection of Ketamine and Xylazine (90 and 10 mg/kg body weight). A drop of 0.3% hypromellose (GenTeal Tears, Alcon, Fort Worth, TX) was applied to the corneas to cushion the contact with the electrode stimulators. Scotopic responses were recorded at stimulus intensities of 0.01, 0.1, 1, 10, and 32 cd*s/m^2^. The rats were then light adapted for 10 min before photopic testing. Photopic responses were recorded sequentially using 10, 32, and 100 cd*s/m^2^ stimuli. Photopic flicker responses were then recorded using 20 cd*s/m^2^ cycled at 9.9 Hz. During testing, body temperature was maintained to 37°C by the heating element built into the Diagnosys Celeris System.

## 3. Results

### 3.1. Retinal Vascularization

The effects of Lrp5 deficiency on the superficial vasculature have been reported in detail in our previous communication (Ubels et al., 2020). For context, two previously unpublished images of the superficial vasculature of wild-type and *Lrp5* knockout rats are provided in Fig. 1. Note that in the wild-type rat, the vasculature extends to the periphery of the retina and is well organized, with abundant capillaries that branch regularly. In contrast, the retinal vasculature of knockout rats is sparse, with reduced branching compared to the wild-type retina. There are large avascular areas where blood vessels do not reach the periphery, and prominent regions with autofluorescent exudates are also present.

**Fig. 1.**
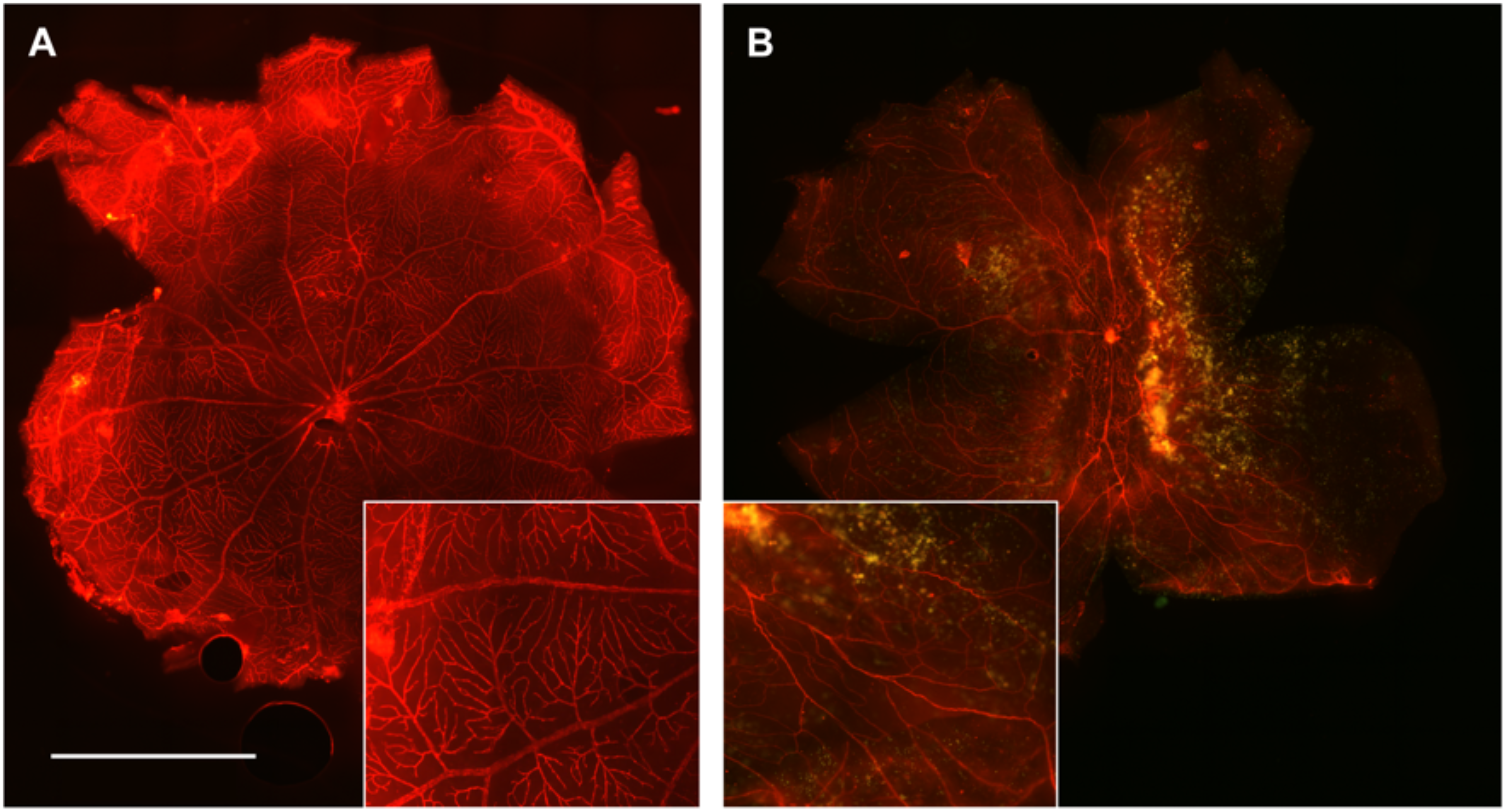
Effect of an 18 bp deletion in the *Lrp5* gene on the superficial vasculature of the 3-month-old rat retina. Flat-mounted retinas of wild-type (A) animals stained with Alexa Fluor 594-conjugated isolectin B_4_ have well-organized blood vessels that branch regularly to the retinal periphery. Vessels in the retina of the *Lrp5* knockout rat (B) are sparse and disorganized with reduced branching. Extensive autofluorescent exudates are present. Insets: Detail of the vasculature magnified 2X from the original. The images are representative of 8 wild-type and 6 knockout rats.

We have now extended the investigation to the deeper vascular plexuses to obtain information on the retinal microvasculature of *Lrp5* knockout adult rats. Fluorescence images of frozen sections of wild-type retinas stained with DAPI and IB_4_ show normal retinal anatomy with three vascular beds present (Fig. 2A). Note that individual nuclei are not clearly defined by DAPI because relatively thick, 12 μm sections were cut to ensure the presence of blood vessels in each section. The superficial plexus is present in the nerve fiber layer of the retina overlying the ganglion cell (GC) layer, the intermediate vessels are visible in the inner plexiform layer (IPL), as is the deep vascular network in the outer plexiform layer (OPL) (Fig. 2B & C). The vascular anatomy is abnormal in the *Lrp5* knockout rat retinas (Fig. 2D & G). Superficial vessels are present, but no intermediate and deep vessels are apparent (Fig. 2E, F, H & I). Retinal layers are also missing and autofluorescent exudates are visible in Fig. 2H. Frozen sections from retinas of rats with a 22 bp deletion were also analyzed, and a comparable vascular phenotype was found with large exudates, as shown in Supplemental Fig. 1.

**Fig. 2.**
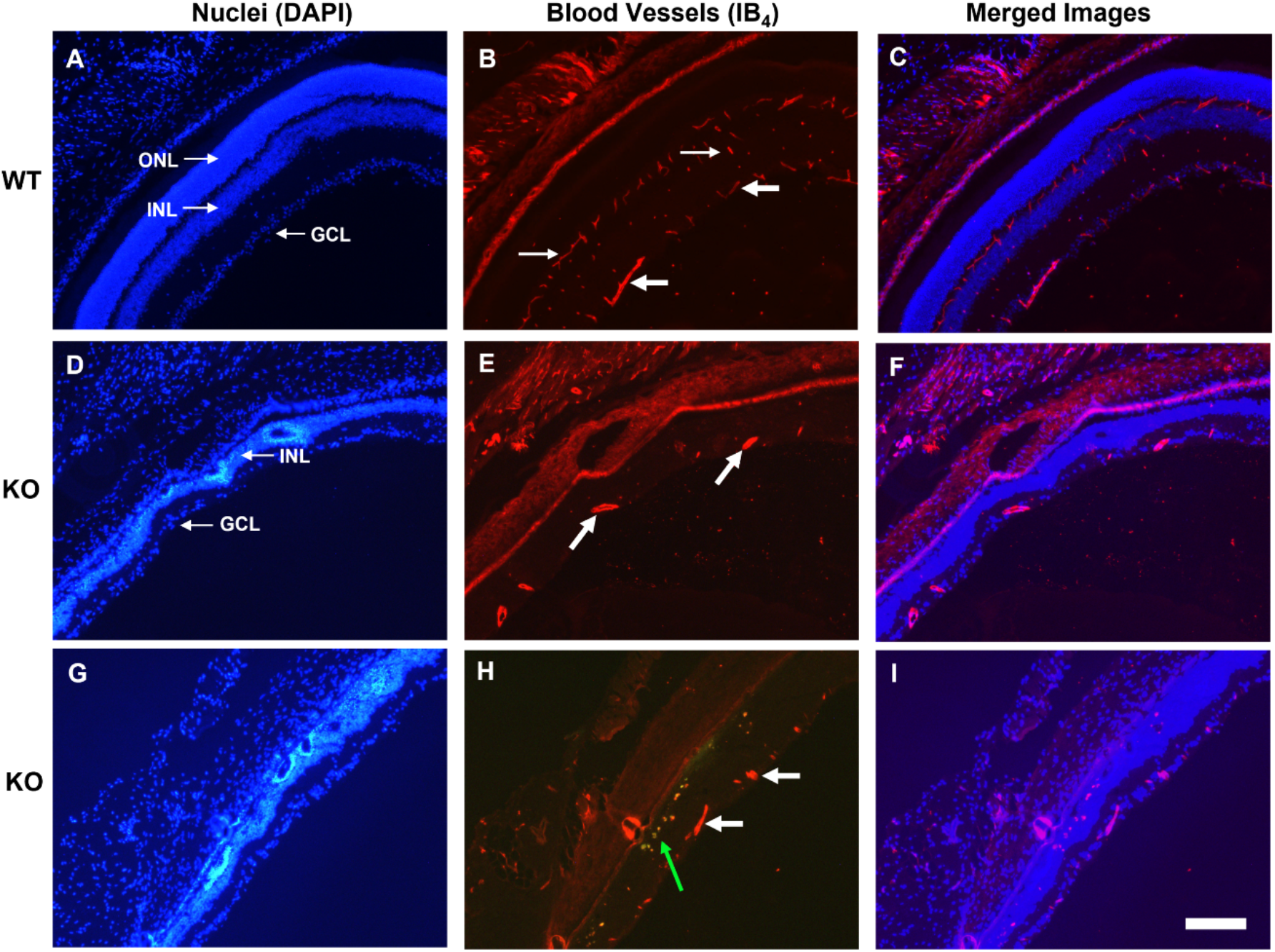
Frozen sections of wild-type (WT) and 18 bp *Lrp5* knockout (KO) retinas showing the retinal vasculature. Images A, D, and G show the retinal layers stained with DAPI. In image A, the outer nuclear layer (ONL), inner nuclear layer (INL), and ganglion cell layer (GCL) are labeled. B, E, and H show the vasculature stained with IB_4_. Images C, F, and I are merged images showing the relationship of the blood vessels to the retinal layers. Wild-type image A shows normal retinal anatomy, while it is evident that outer retinal layers are absent in KO retinas (D and G). Superficial retinal blood vessels are present in both WT and KO retinas (B, E, and H, large arrows). The WT retina has normal intermediate and deep vascular plexuses (B, small arrows, and C). In contrast, the intermediate and deep plexuses are absent from the KO retinas (E, H, F, and I). Autofluorescent exudates are present in KO retinas (H, green arrow). These exudates appear pink in the merged image (I) and should not be mistaken for blood vessels. The images are representative of 3 wild type and 3 knockout rats. Scale bar = 100 μm

### 3.2. Claudin-5 Expression

In wild-type rats, the superficial vessels and the intermediate and deep vascular plexuses all stain for claudin-5 (Fig. 3A). The pattern of staining corresponds to the pattern of staining of these vessels with isolectin B_4_ (Fig 2C). No staining for claudin-5 is present in the *Lrp5* knockout retinas Fig. 3B & C). Note that although superficial vessels are present in knockout retinas (Fig. 2E & H), these vessels do not stain for claudin-5 in knockout retinas. Large fluorescent structures are present in the knockout retinas, but they do not represent claudin-5. The same autofluorescent inclusions are visible in knockout retinas that were not exposed to the Alexafluor 488-conjugated claudin-5 antibody (Fig. 3D), and thus it is concluded that they are exudates.

**Fig. 3.**
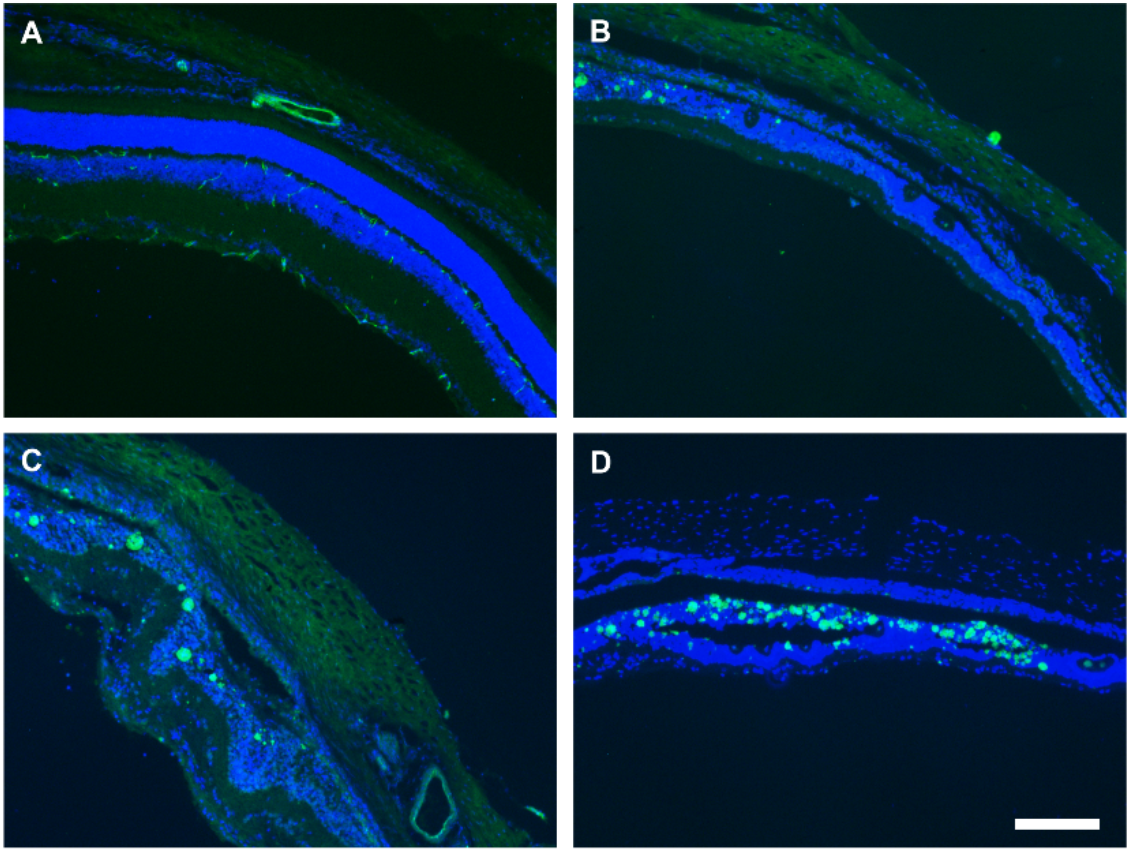
Frozen sections of wild-type and knockout retinas stained for claudin-5. Claudin-5 is expressed in superficial, intermediate, and deep vessels of the wild-type rat (A). There is no staining for claudin-5 in knockout retinas (B & C). The large fluorescent inclusions visible in the knockout retinas are also present in knockout retinas that were not exposed to anti-claudin-5 (D) and represent autofluorescent exudates. The images are representative of 3 wild-type and 3 knockout rats. Scale bar = 100 μm

### 3.3. Retinal Histology

Since frozen sections showed compromised retinal anatomy with missing layers in knockout rats, the retinal structure was investigated in more detail by histology of paraffin sections with H & E staining. Fig. 4A shows the normal anatomy of a wild-type retina. *Lrp5* knockout retinas do not have photoreceptors (Figs. 4B and C), and exudates are present in the inner nuclear layer (Fig. 4C). The thickness of the inner nuclear layer is reduced from 28.0 ± 4.9 μm in wildtype rats to 20.2 ± 2.9 μm in knock out animals (mean ± SD, significant difference, t-test, p ≤ 0.05, n=11). Similarly, the thickness of the inner plexiform layer was reduced from 46.5 ± 10.3 μm to 18.4 ± 3.6 μm (n=12).

**Fig. 4.**
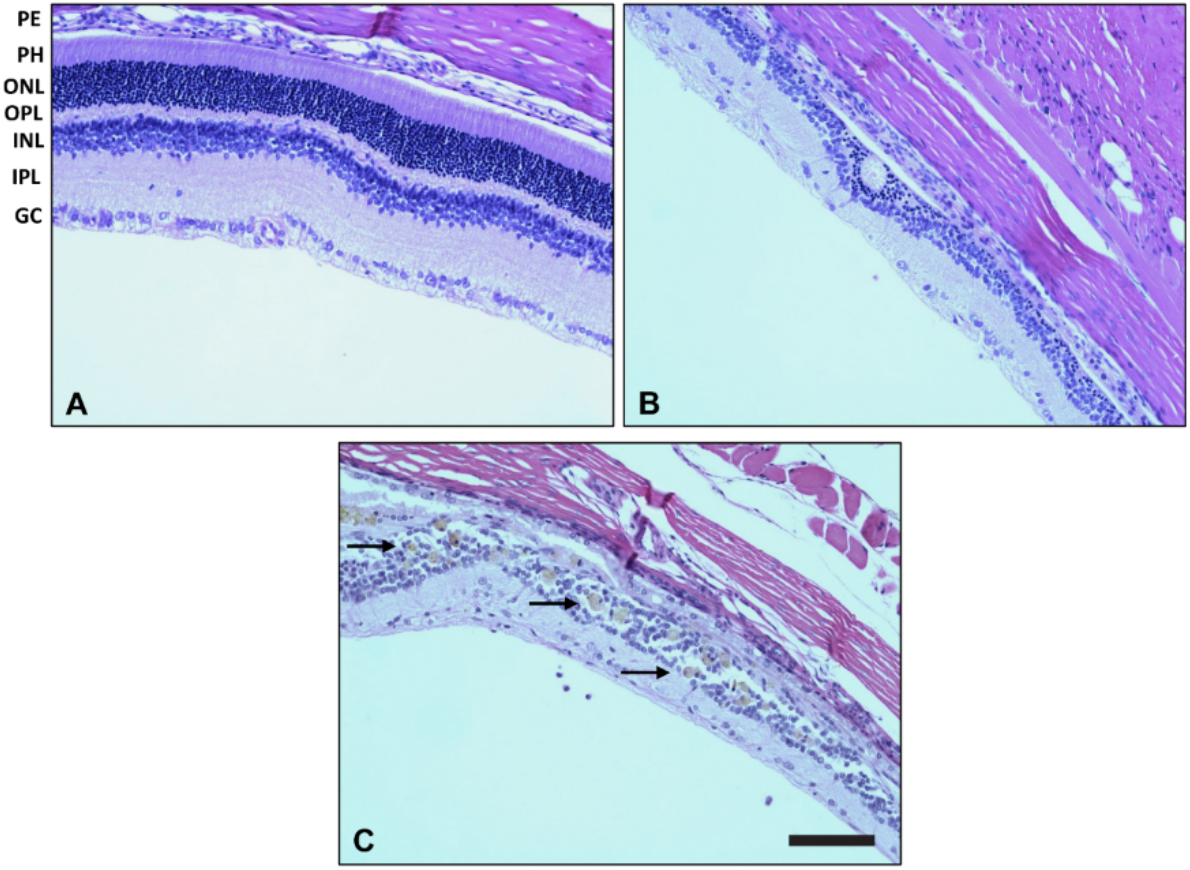
Histologic sections of rat retinas stained with H&E. The wild-type image (A) shows the normal anatomy of the retinal layers. Retinas of knockout animals do not have photoreceptors or an outer plexiform layer and contain large exudates (B & C). The ganglion cell layer is present but sparse and disorganized. The inner nuclear and inner plexiform layers are thinner than in the wild-type retina. The images are representative of 2 wild-type and 3 knockout rats. Scale bar = 100 μm

### 3.4. In Vivo Retinal Architecture

Fundus images of wild-type retinas revealed typical vascular anatomy, with 10 - 11 large vessels radiating from the optic nerve head. In contrast, fundus images of knockout animals show fewer large vessels and a disorganized vasculature (Fig. 5), which agrees with our previously reported quantitative data on the numbers of superficial vessels in knock out rats (Ubels et al., 2020). The image is hazy suggesting presence of a pre-retinal membrane which we have previously observed during dissection of Lrp5-deficient retinas for flat mounts (unpublished data).

**Fig. 5.**
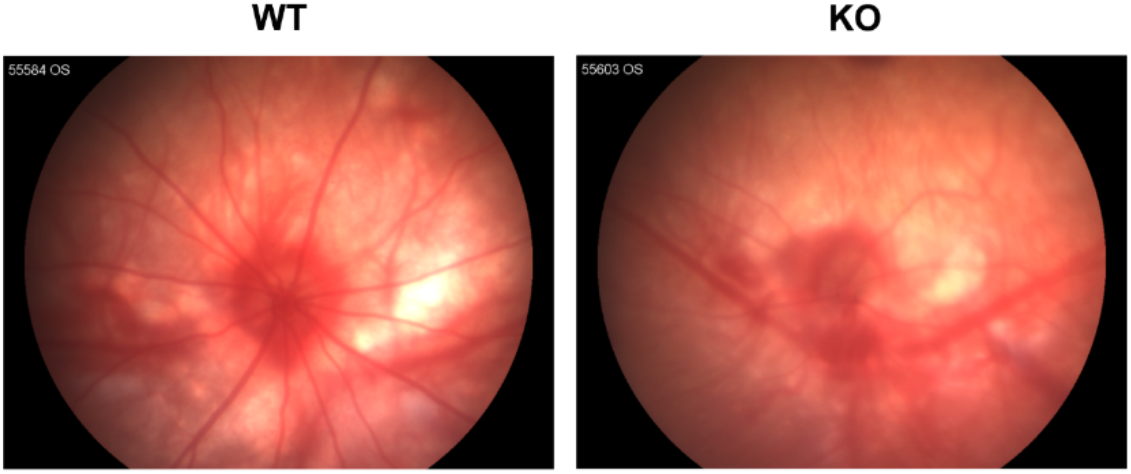
Fundus photography of wild-type (WT) and knockout (KO) rat retinas. WT retinas have normal retinal vasculature. KO retinas have a sparse vasculature. The images are representative of 5 wild-type and 4 knockout rats.

In contrast to the normal structure of wild-type retinas, OCT of knockout rats shows thinning of the retina with missing layers (Fig. 6). In agreement with fluorescence and light microscopic images (Figs.1–4), the retina of knockout rats has abundant hyperreflective foci, consistent with the presence of exudates (Fig. 6B–D). Prominent folds in the retina were also present (Fig. 6D). Retinal thickness was measured 2 mm from the optic nerve head. As shown in Figure 7, wild-type rats had a thickness of 187.2 ± 3.4 μm. In contrast, OCT of *Lrp5* knockout rats shows a significantly reduced thickness of 78.2 ± 5.5 μm, with photoreceptors and the outer nuclear layer missing. In agreement with the thin inner nuclear layer of knockout rats seen in Fig. 4, the inner nuclear layer is not clearly discernable on OCT (Fig. 6B–D).

**Fig. 6.**
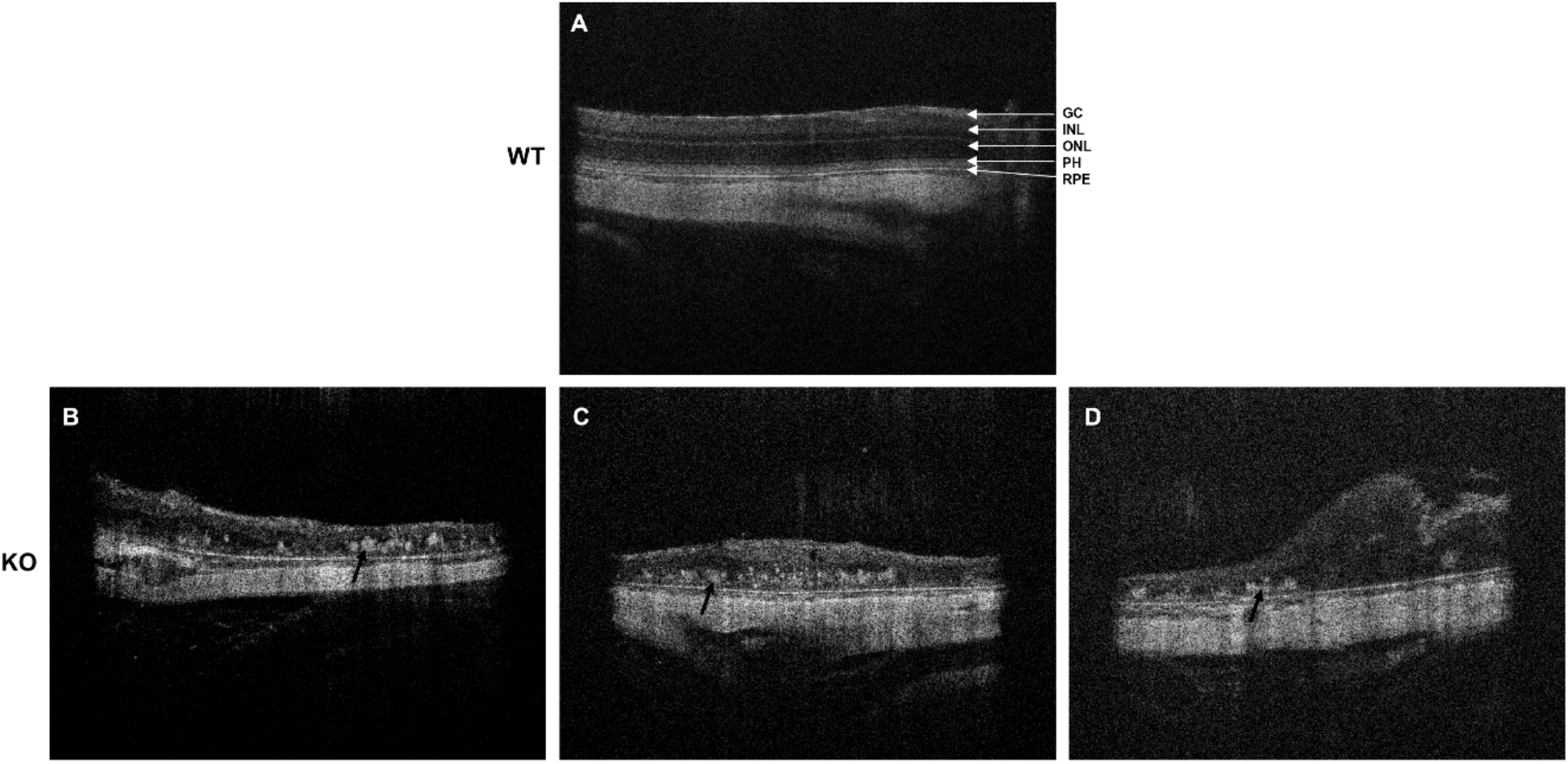
OCT of the retina 2 mm peripheral to the optic nerve head. On the wild-type (WT) image (A), the retinal pigment epithelium (PE), photoreceptor layer (PH), outer nuclear layer (ONL), and inner nuclear layer (INL) and ganglion cell layer (GC) are labeled The PH layer and ONL are missing, and the INL is not clearly discernable in KO retinas (B). Exudates are present in the knockout retinas (arrows) and there is a large retinal fold in image C. The images are representative of 5 wild-type and 4 knockout rats.

**Fig. 7.**
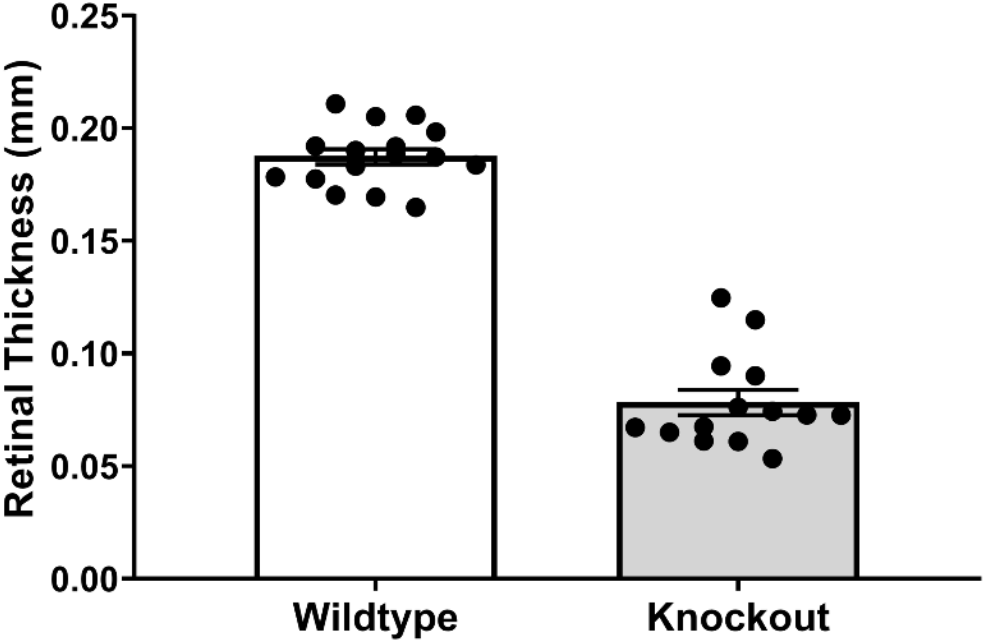
Retinal thickness measured 2 mm from the optic nerve head. The retina of knockout rats is significantly thinner than the wild-type retina. (*significant difference, t-test, p ≤ 0.05; WT, n = 16; KO, n = 14)

### 3.5. Vascular Permeability

The exudates in knockout retinas suggest that deficiency of Lrp5 results in a defective BRB. Measurement of vascular permeability with Evans blue dye confirmed this. Retinas from wild-type rats had an average of 11.6 ± 2.4 μl/g/h (n = 14) of accumulated Evans blue dye. In contrast, the dye accumulation was increased 3.6-fold in *Lrp5* knockout rats to 41.9 ± 7.9 μl/g/h (n = 10; Fig. 8). It should be noted that the increased Evan’s Blue dye occurred with a significant decrease in capillary surface, as previously reported (Ubels et al., 2020), indicating a much higher degree of permeability than the observed 3.6-fold accumulation of dye.

**Fig. 8.**
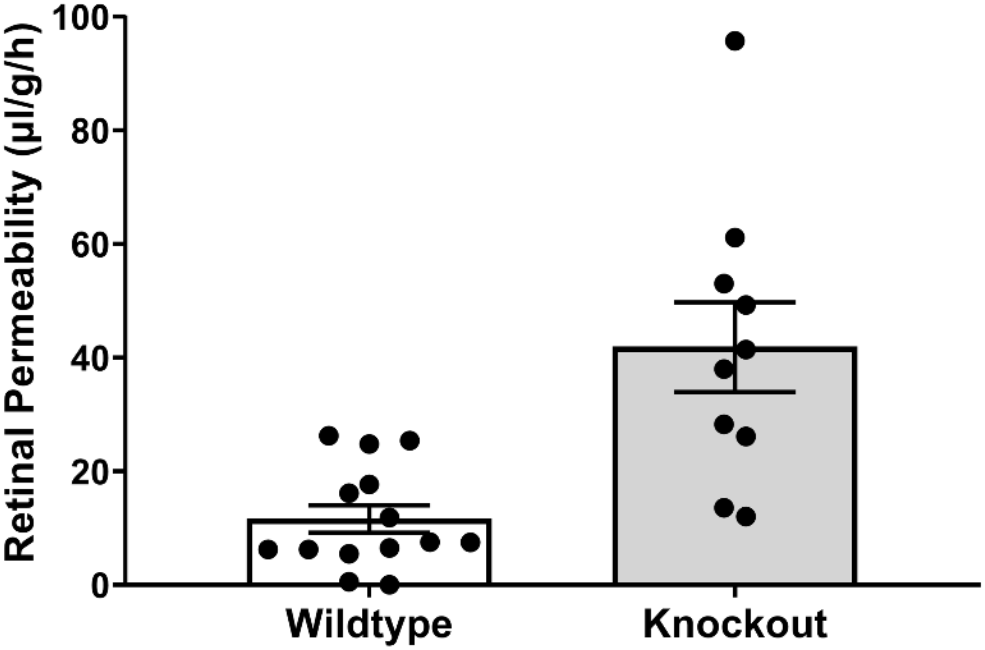
Permeability of retinal vasculature to Evans blue dye. Permeability of vessels of knockout rats is significantly greater than that of wild-type animals. (*significant difference, t-test, p ≤ 0.05, WT, n = 14; KO, n = 10)

### 3.6. Retinal Function

Electroretinograms (ERG) of wild-type animals were typical for albino rats, with prominent scotopic and photopic a- and b-waves (Fig. 9A; Suppl. Figs. 2 and 3). Consistent with the profound abnormalities of the retinal vasculature and neural retina, *Lrp5* gene deletion resulted in an essentially absent electroretinographic response to light (Fig. 9B; Suppl. Figs. 2 and 3).

**Fig. 9.**
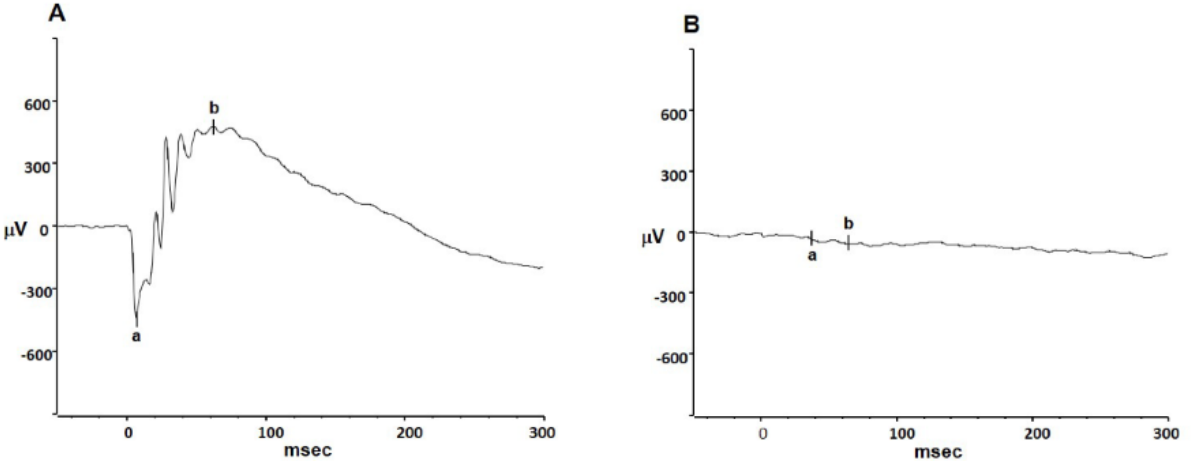
Representative scotopic electroretinograms of wild-type (A) and knockout (B) rats at a stimulus intensity of 32 cd.s/m^2^ at which the WT response reached its maximum. The *Lrp5*-deficient retina is unresponsive to light.

## 4. Discussion

This study shows that deficiency of Lrp5 causes profound structural and functional abnormalities in the rat retina. The superficial and inner vascular plexuses do not develop normally, exudates are present, and the blood-retinal barrier is highly permeable. Histology and in vivo imaging show that these changes are accompanied by abnormal organization of the retinal layers, leading to an impaired response to light. Taken together, these changes indicate that this new rat model should be useful for the study of FEVR and related hypovascular retinal diseases (Chen et al., 2012; Huang et al., 2016; Wang et al., 2012; Wang et al., 2016; Wang et al., 2019; Ye et al., 2009; Xia et al., 2010; Zhou et al., 2014).

Our CRISPR-Cas9 model confirms and extends observations in mice. The retina of both mice and rats has a hypovascular phenotype. The superficial vasculature is present but develops abnormally and the intermediate and deep plexuses are absent. Most mouse models cited above have been studied in the early post-natal period at P7 – P17, focusing on the delayed growth of the superficial vasculature toward the retinal periphery. The sparse, disorganized vasculature is similar in adult, P30 mice (Huang, et al. 2016) and the 3-month rats used in the present study. Our study of adult rats has the advantage of demonstrating the more advanced disease caused by Lrp5 deficiency. A notable difference between the rat and mouse models is the presence of large autofluorescent exudates in the Lrp5-deficient rat retina which appear as brown deposits in light micrographs of histological sections and as hyper-reflective foci on OCT. These exudates have not been reported in *Lrp5* knockout mice either in fluorescence micrographs or on OCT Chen et al., 2012; Huang et al., 2016; Wang et al., 2016).

The presence of these exudates provides evidence of the strength of the rat as a model of human exudative disease (Gilmour, 2015; Kashani et al., 2014; Madan and Penn, 2003; Wang et al 2019), since similar exudates are visible on fundus photos of humans with FEVR (Kashani et al., 2014; Wang et al., 2019) and OPPG (Maltese et al., 2017). Notably, the exudates detected by OCT in the rat are essentially identical in appearance to exudates in the retinas of FEVR patients examined by OCT (Yonakawa et al., 2015). Such exudates were not present on OCT images of Lrp5 knockout mouse retinas (Wang et al., 2016).

Fluorescein angiography and FITC-dextran perfusion in P21 and P30 mice, respectively, have shown that the BRB is defective in the absence of Lrp5 (Huang et al., 2016; Xia et al., 2008). We have now shown directly that the BRB in rats is defective by measuring permeability to Evan’s blue dye, indicating that development of junctional complexes is compromised in the absence of Lrp5. Claudin-5 is an essential component of junctional complexes in retina vessels (Diaz-Coranguez, et al. 2017). In Lrp5-deficient mice, the claudin-5 gene is down-regulated (Chen et al., 2012), and we have now shown that the claudin-5 protein is not expressed in the superficial vessels of *Lrp5* knockout rats.

A novel finding of the present study is that, in addition to vascular pathology, Lrp5-deficient rats have profound abnormalities in the structure of the neural retina. Retinal thickness was reduced by 50% as measured by OCT, and histology confirmed that this reduction in thickness was due to the absence of photoreceptors and the outer plexiform layer and a reduction in thickness of the inner nuclear and inner plexiform layers. In contrast, OCT of Lrp5 knockout mice at P30 demonstrated minimal changes in retinal thickness (Wang et al., 2016), and all retinal layers were present. OCT of the knockout rats shows features similar to human FEVR. Yonekawa et al. (2015) have conducted the most detailed OCT study of FEVR patients to date. They demonstrated retinal folds, thinning, and loss of outer layers. The data suggest that the rat model is an improvement over the mouse models with respect to retinal structure.

Consistent with both microscopic imaging and OCT that revealed the absence of the photoreceptor and intraneuronal layers in Lrp5 knockout rats, the knockout retina is unresponsive to light. This finding differs from mouse Fzd4 and Lrp5 knockout models in which the ERG shows a negative deflection, consistent with the presence of photoreceptors, but no b-wave. This is consistent with milder pathology in mice with a defective Wnt signaling pathway (Ye et al., 2009; Wang et al., 2016). Together, these results in rats and mice agree with a report by Yukari et al. (2016) who showed that while the ERG is normal in FEVR patients with mild vasculopathy and no retinal folds, there is a progressive reduction in ERG amplitude as vascular and structural changes become more severe. This eventually results in complete loss of the response to light in patients with abnormalities in all major retinal vessels. The loss of visual function in the Lrp5-deficient rat appears to model the effect of severe FEVR on the response of the retina to light.

There is a need for the development of therapies for retinal exudative diseases including FEVR, OPPG, Norrie disease, retinopathy of prematurity, and diabetic retinopathy, all of which involve pathological neovascularization coupled with incomplete barriergenesis related to defects in Wnt signaling; however, there have been some studies of treatment of these conditions in animal models. In Lrp5-deficient mice, lithium chloride injected intraperitoneally to systemically activate the Wnt/β-catenin signaling pathway partially promoted the angiogenesis of the deeper vasculature Wang et al. (2016). Treatment with norrin has a positive effect on the BRB in diabetic rats and promotes revascularization of the hypoxic area of the retina in a mouse model of OIR (Diaz-Coranguez et al., 2020; Ohlmann et al., 2005; Ohlmann et al., 2010). Clearly, additional research on treatment of hypovascular retinal disease is required.

CRISPR-Cas9 gene-editing technology has resulted in the reemergence of the rat as a predictable organism for studying the physiology and pathology of human disease, with the advantage that rats have larger organs than mice, providing easier dissection and larger tissue samples (Holmberg et al, 2017; Neff, 2019). Our use of CRISPR-Cas9 in the rat demonstrates the feasibility of using this technology for the study of genetic diseases of the retina. We propose that the model presented in this report will be useful for further studies of hypovascular, exudative retinal vascular diseases that involve the Wnt and norrin signaling pathways and the development of treatments for these conditions. These rats should also be useful for studies beyond the scope of the present study, such as investigation of the outer BRB in the choroidal vasculature and the mechanisms by which Lrp5-deficiency impairs development of photoreceptors and the neural retina. To make the *Lrp5* knockout rats available to investigators who wish to do more detailed studies of Wnt signaling in the neurovascular unit of the retina, the rat strains that we have produced have been submitted to the Rat Resource and Research Center at the University of Missouri.

## Acknowledgments

This work was funded by the Van Andel Institute, the Calvin Fund for Eye Research, and NIH Vision Research Core Grant EY007003 at the Kellogg Eye Institute. The following members of the VAI Vivarium and Transgenics Core provided technical assistance: Bryn Eagleson, Adam Rapp, Nicholas Getz, Audra Guikema, Tristan Kempston, Malista Powers, and Tina Schumaker.

The rat strains have been deposited with the Rat Resource & Research Center (RRRC) at the University of Missouri, Columbia, MO (https://www.rrrc.us) for public distribution. The strains have the numbers: 18 bp deletion, RRRC #867; 22 bp deletion, RRRC #880.

## Supplementary Data

Ubels et al., Structure and Function of the Retina of Low-density Lipoprotein Receptor-related Protein 5 (Lrp5)-deficient Rats

**SM Fig. 1.**
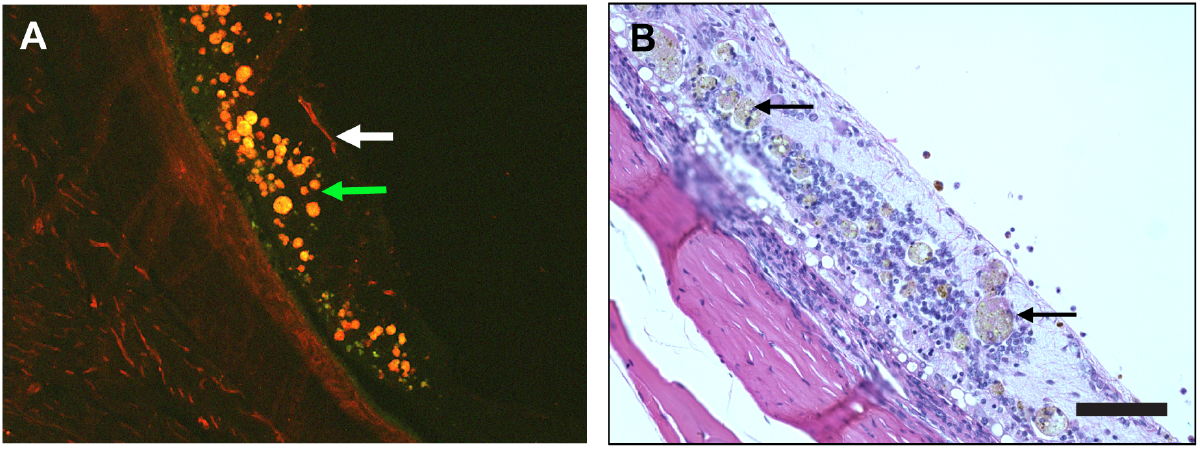
Retinas of rats with a 22 bp deletion in Lrp5 were observed in our previous report (Ubels et al., 2020) to have more extensive exudates than rats with the 18 bp deletion. A. Frozen section from a 22 bp deletion rat stained with IB_4_. Large exudates are present (green arrow). The white arrow indicates a superficial vessel, but the intermediate and deep plexuses are absent. C. Paraffin section stained with H & E. Note missing photoreceptors and large exudates (arrows). Scale bar = 100 ⍰m

**SM Fig. 2.**
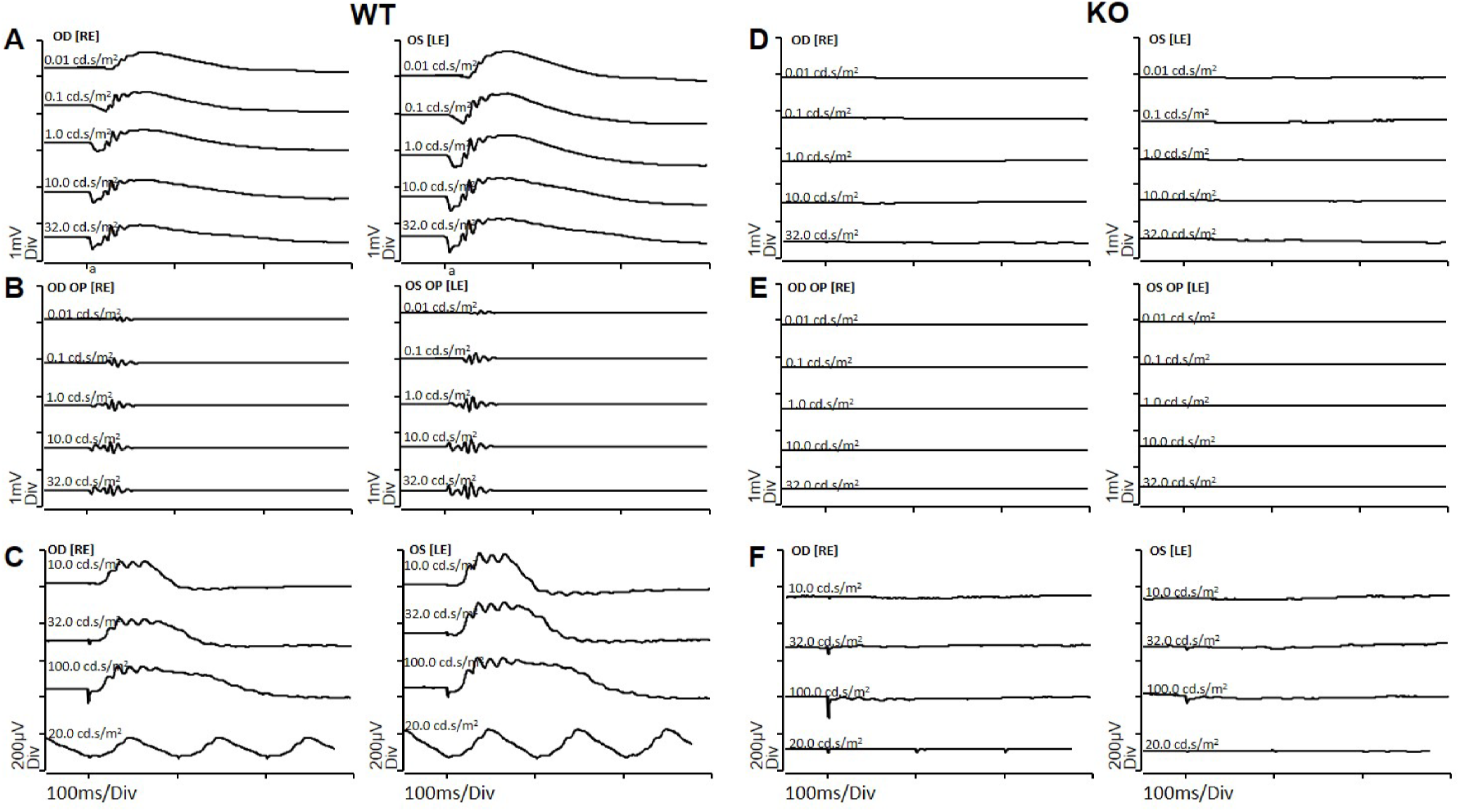
Representative ERG waveforms of wild-type and Lrp5 knockout rats. Representative scotopic responses (A & D) and their oscillatory potentials (B & E) were recorded at stimulus intensities of 0.01, 0.1, 1, 10, and 32 cd*s/m2. Rats were then light-adapted for 10 minutes before photopic testing. Photopic responses were recorded using 10, 32, and 100 cd*s/m2 stimuli plus a flicker stimulus of 20 cd*s/m2 cycled at 9.9 Hz (C & F).

**SM Fig. 3.**
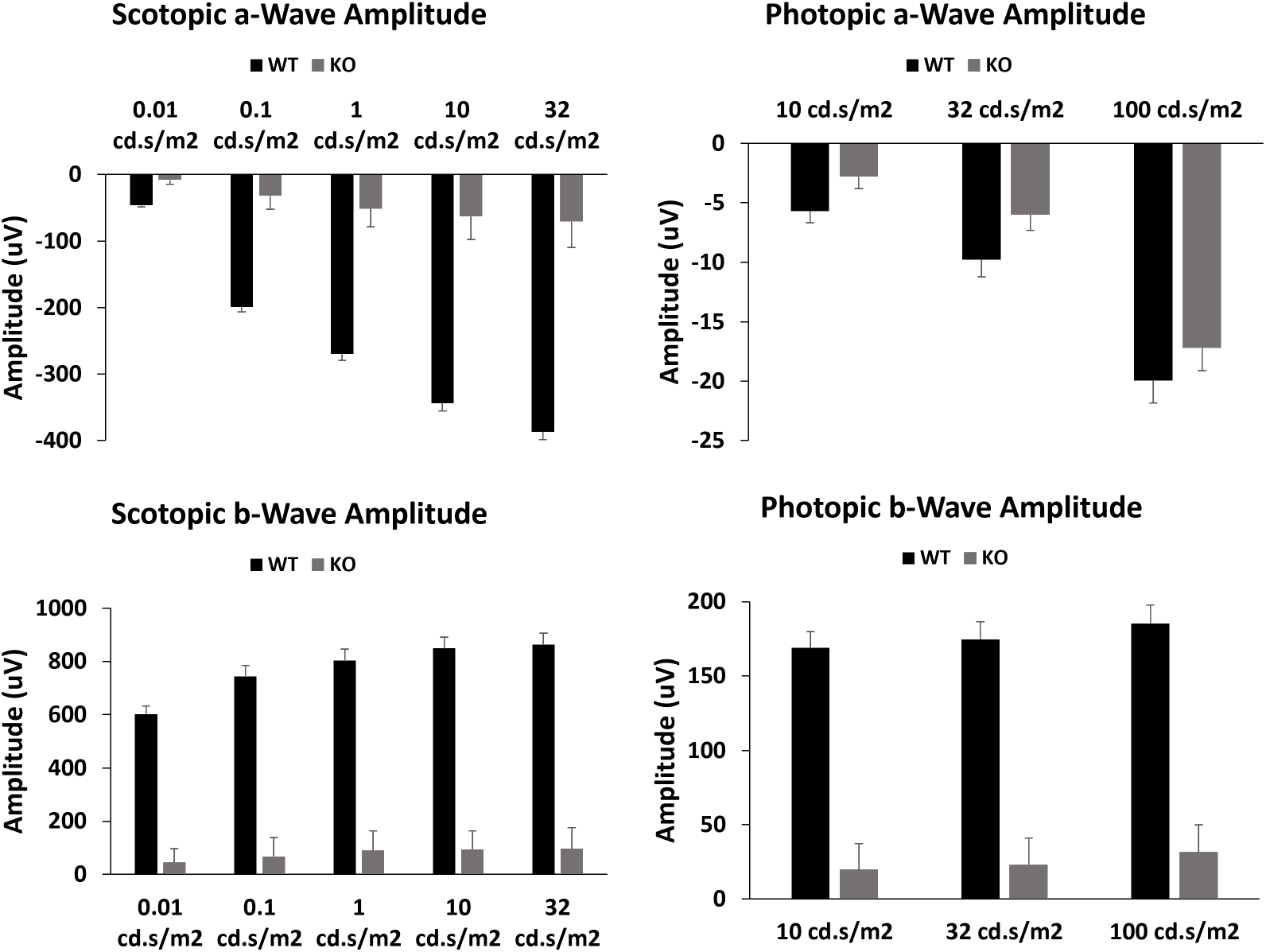
ERG amplitudes of wild-type and Lrp5 knockout rats. (mean + SD, WT: n=5; KO: n=4)

